# ATM-Mediated Translocation of RanBPM Regulates DNA Damage Response by Stabilizing p21 in Non-Small Cell Lung Cancer Cells

**DOI:** 10.1101/2022.12.20.520942

**Authors:** Tanggang Deng, Lin Xie, Yugang Xiao, Zhenbin Zhang, Yuchong Peng, Linglong Yin, Yongming Fu, Xiong Li

## Abstract

Non-small cell lung cancer (NSCLC) is the leading cause of cancer-induced deaths around the world, and platinum-based chemotherapy remains a standard-of-care for most patients with advanced NSCLC. DNA damage response (DDR) induced by platinum or Etoposide activated a panel of cell cycle-regulatory proteins including p21 through p53 pathway. In this present study, we found that the level of p21 or RanBPM is lower in NSCLC than non-malignant tissues and has a highly positive correlation, which is negatively correlated with the survival of patients. We further revealed that RanBPM protein physically interacts with p21, RanBPM deubiquitinates p21 by recruiting a deubiquitinase USP11 to maintain protein stability of p21. Furthermore, RanBPM regulates DNA damage response (DDR) in a p21-dependent manner, and DNA damage promotes the translocation of RanBPM into the nucleus and regulates p21 protein stability through ATM-mediated pathways. We for the first time revealed a novel mechanism of p21 protein stability regulated by RanBPM, and the novel roles of RanBPM in the regulation of DDR.

## Introduction

Lung cancer is a frequently diagnosed malignancy with the second morbidity and the highest mortality worldwide. According to the latest Global Cancer Statistics 2020, lung cancer accounts for 11.4% of the total new cases, and for 18% of the total cancer deaths[1]. Non-Small Cell Lung Cancer (NSCLC) is the most common type of lung cancer, accounting for 80–85% of all cases[2]. Based on the histological features, NSCLC is further divided into lung adenocarcinomas, squamous cell carcinoma and large cell carcinoma[3]. Platinum-based chemotherapy is the standard-of-care for NSCLC patients before or after surgery, or alongside radiotherapy, or for the advanced NSCLC patients[4]. Chemotherapy-caused DNA damage activates a series of DNA damage response (DDR), including the activation of ATM/ATR, and then triggers the downstream cell processes, such as cell cycle arrest, DNA damage repair, cell senescence or apoptosis[5]. However, cancer cells will develop drug-resistance after a period of treatment. Therefore, it is urgently required to reveal the molecular mechanism of drug resistance of NSCLC, identify the predictive diagnostic biomarkers, and develop the new therapies.

RanBPM, known as RanBP9, is a ubiquitous, evolutionarly conserved scaffold protein, which is localized in both cytoplasm and nucleus[6]. Previous studies have demonstrated that RanBPM interacts with multiple proteins that involved in various cellular processes, such as cell adhesion and migration, DDR and signal transduction[7–9]. RanBPM-deficient mice show early perinatal lethality and sterility[10, 11]. RanBPM has also been identified as a critical regulator of protein stabilization, and regulates diverse biological functions through protein interaction. RanBPM sustains the protein stabilization of Mgl-1, and promotes its tumor suppressor activity [12]. However, the detailed mechanism underlying the effect of RanBPM on DDR of NSCLC cells remains unclear.

As a broadly acting cyclin-dependent kinase inhibitor, p21 plays a critical roles of tumor suppressor. The dysregulation of p21 has been reported in multiple human cancers [13]. In normal physiological conditions, p21 protein is rapidly degraded by the ubiquitin-proteasome pathway [14, 15]. However, in response to diverse cellular stress, such as DNA damage, the protein levels of p21 rapidly elevated, which results in cell cycle arrest, apoptosis or cellular senescence[16]. Recent studies have revealed that USP11 deubiquitinates and stabilizes p21 protein under physiological conditions, as well as in response to DNA damage[17].

In this present study, we at the first time reported a new molecular mechanism by which RanBPM sustains the protein stability of p21 in a USP11-dependent manner. In particular, we found that DNA damage promoted the ATM-mediated nuclear translocation of RanBPM protein, and colocalized and physically interacted with p21. Taken together, RanBPM has been identified as a novel regulator of p21 protein stability, and plays a critical role in the regulation of DDR.

## Materials and Methods

### Cell Cultures

HEK293T, A549 and H1299 cells were purchased from American Type Culture Collection (ATCC, Manassas, VA). A549 and H1299 cells were cultured in RPMI 1640 medium (Gibco, ThermoFisher Scientific, Friendship, ME, USA) supplemented with 10% fetal bovine serum (FBS, Thermo Fisher Scientific, Waltham, MA, USA) plus 5 mM glutamine, penicillin G (100 U mL^-1^) and streptomycin (100 μg mL^-1^) at 37°C under 5% CO2. HEK293T were maintained in Dulbecco’s Modified Eagle Medium (DMEM) supplemented with 10% FBS and 1% penicillin/streptomycin. The experiment protocol was approved by the Ethics Committee of the Guangdong Pharmaceutical University.

### Antibodies and chemicals

Specific antibodies against p53 (DO-1, sc-126), p15 (D-12, sc-271791), p19 (SPM429, sc-65594) and RanBPM (sc-271727) antibodies were purchased from Santa Cruz Biotechnology (Dallas, Texas, USA), and p18 (#2896), p21 (#2947), p57 (#2557) and RanBPM (#14638) antibody were purchased from Cell Signaling Technology (Danvers, MA, USA). Anti-p16 (A301-267A) was purchased from Bethyl Laboratories (Montgomery, TX, USA). Anti-p27 (AF1669), anti-ATM (AF1399) and anti-phospho-ATM (Ser1981) were purchased from Beyotime Biotechnology (Shanghai, China). Anti-USP11 (ab109232) and anti-RanBPM (ab205954) antibodies were purchased from Abcam (Shanghai, China). Anti-GAPDH (AT0002) antibodies were purchased from CMCTAG (Milwaukee, WI, USA). Anti-Flag (cat. M185-3L), anti-Myc (cat. M192-3) and anti-HA (cat. M180-3) antibodies were purchased from Medical & Biological Laboratories CO., LTD. (Minato-ku, Tokyo, Japan). Etoposide (VP-16, S1225) and KU-55933 (S1092) were purchased from Selleck Chemicals (Shanghai, China), and Cycloheximide (sc-3508B) and MG132 (C2211) were purchased from Sigma (Shanghai, China). Doxorubicin (SC0159) was purchased from Beyotime Biotechnology (Shanghai, China).

### Western Blotting and Immunoprecipitation

The procedures of western blotting and Immunoprecipitation assay were performed as previously described[17].

### GST Pulldown Assays

GST fusion proteins and His fusion proteins were produced following standard protocol. For in vitro binding assays, bacterially expressed GST-p21 bound to glutathione Sepharose beads (Thermo Scientific, 16100) were incubated with His-RanBPM. After washing, the bound proteins were separated by SDS-PAGE and immunoblotted with indicated antibodies.

### Protein Half-Life Assays

Cells were transfected with the indicated siRNA for 48 h, or transfected with specified plasmids for 24 h followed by treatment with/without cycloheximide (50 μg·mL^-1^) for various periods of time, washed with PBS, and lysed in RIPA buffer containing a protease inhibitor cocktail. The protein levels were assessed using Western blot analysis.

### Real-time PCR

Total RNA was extracted using Trizol (Takara Bio, Otsu, Japan). RNA (1 μg) was reverse-transcribed in a 20 μL reaction using RevertAid First Strand cDNA Synthesis Kit (Thermo Scientific, #K1622). After reverse transcription at 42°C for 60 min, then 42°C for 15 min and inactivation by incubating samples at 70°C for 5 min, the RT reaction was diluted. cDNA was used for RT-PCR or real-time PCR assay. Primer sequences were as follows:

RanBPM-F: GGTGATGTCATTGGCTGTTG

RanBPM-R: AATTTGGCGGTAGGTCAGTG

GAPDH-F: AAGGTGAAGGTCGGAGTCAA

GAPDH-R: AATGAAGGGGTCATTGATGG

p21-F: ATTAGCAGCGGAACAAGGAGTCAGACAT

p21-R: CTGTGAAAGACACAGAACAGTACAGGGT

### RNA interference

The sequences of the RanBPM siRNAs have been previously reported:

siUSP11#1: 5’-AAUGAGAAUCAGAUCGAGUCC-3’

siUSP11#2: 5’-AAGGCAGCCUAUGUCCUCUUC-3’

siRanBPM#1: 5’-GGCCACACAAUGUCUAGGA-3’,

siRanBPM#2: 5’-GGAAUUGGAUCCUGCGCAU-3’, and the sequence of the control siRNA is 5’-UUCUCCGAACGUGUCACGUUUC-3’. All these siRNAs were synthesized by Shanghai GenePharma. siRNA transfection was performed in cells using GeneMut siRNA transfection reagent (SignaGen Laboratories cat. SL100568) (Rockville, MD, USA). The experimental procedure followed the protocol provided by the manufacturers.

### *In vivo* ubiquitylation assay

HEK293T cells were transfected with the indicated siRNAs, or transfected with the indicated plasmids for 24 h followed by treatment with 20 μM MG132 for 6 h, washed with PBS, and lysed in RIPA buffer containing a protease inhibitor cocktail. The lysates were transferred into a 1.5 mL tube and placed on a hot plate immediately to boil for 10 min. Then the lysates were incubated with anti-His antibody overnight, followed by treatment with protein A/G beads for an additional 2 h at 4°C. After three washes with PBS buffer containing 1%o Tween-20 (PBST), ubiquitinated p21 was analyzed using immunoblotting with anti-HA antibody.

### Clonogenic survival assay

Cells were treated with the indicated drugs, then collected, counted, diluted, then finally seeded into 6-well plate at an appropriate density for each treatment. After 10 to 14 days of incubation, the colonies were fixed and stained with crystal violet for 30 minutes. The colonies (≥0.3mm) with at least 50 cells were counted as survivors under a stereomicroscope by ImageJ software. Data are representative of three independent experiments.

### Immunohistochemical staining analysis

Formalin-fixed, paraffin-embedded samples were sectioned at 5 μM. Sections were treated with antigen retrieval buffer. Specifically, incubation with antibodies against RanBPM (1:50 dilution; Santa Cruz, USA), p21 (1:100 dilution; CST) was carried out overnight at room temperature. Slides were incubated in secondary antibody. The protein levels of RanBPM and p21 in the tumor specimens from NSCLC patients were reviewed and scored under a light microscope. Each specimen was quantified and assigned a score based on the intensity of the membrane, cytoplasmic, and/or nucleic staining by a visual grading system (0-3) (grade 0, no staining; grade 1, weak staining; grade 2, moderate staining, grade 3, strong staining) and the extent of stained cells (0% =0, 1-24% =1, 25-49% =2, 50-74%=3, 75-100%=4). The final immunoreactive score ranges from 0 (no staining) to 12 (strong staining).

### Survival analysis

The effects of RanBPM genes on the survival of patients with NSCLC were analysed using Kaplan–Meier Plotter (http://kmplot.com/analysis/)[26], which scontained gene expression data and survival information of 2,434 clinical NSCLC patients.

### Statistical analysis

All data were analyzed by GraphPad Prism 9.0 (GraphPad Software, San Diego, CA, USA; RRID:SCR_002798) or FlowJo software (FlowJo, RRID:SCR_008520). Results are presented as mean ± SEM or mean ± SD as indicated. The statistical difference between two samples was analyzed by Students t test. One-way ANOVA was used to analyze the statistical difference of multiple groups. *P < 0.05 and **P < 0.01. P < 0.05 was considered as statistically significant.

## Results

### RanBPM is positively correlated with p21, and negatively correlated with the survival of NSCLC patients

To explore the roles of RanBPM in the development of NSCLC, we first analyzed the correlation of the mRNA levels of RanBPM with the survival of NSCLC patients by using publicly available datasets (http://kmplot.com/analysis/). The data of Kaplan-Meier curve and log rank test revealed that the mRNA levels of RanBPM was positively correlated with the overall survival (OS), progression-free survival (FP), and post-progression survival (PPS) of NSCLC patients (p < 0.05, Figure 1A). Furthermore, we performed immunohistochemical (IHC) staining using antibodies against RanBPM on a tumor tissue microarray consisting of 52 NSCLC patient specimens. NSCLC tumor specimens express much lower RanBPM levels than the normal tissues adjacent to the tumor (Figure 1B–C). It was reported that RanBPM knockdown induced cell cycle arrest at S-phase [9]. However, the detailed mechanism remains unclear. We analyzed the correlation of RanBPM (RANBP9) with cyclin-dependent kinases inhibitors (CKIs) by using Gene Expression Profiling Interactive Analysis (GEPIA) web server (Figure 1D and Figure S1A). The expression levels of RanBPM is positively correlated with p21(CDKN1A) in NSCLC tumor samples (Figure 1D). RanBPM knockdown specifically decreased p21 level (Figure S1B). We further validated the correlation of RanBPM with p21 protein levels in NSCLC tumor specimens by IHC analysis (Figure 1E–F). These data indicated that the expression of RanBPM is positively correlated with p21 in the tumor specimens of NSCLC patients.

**Figure 1.**
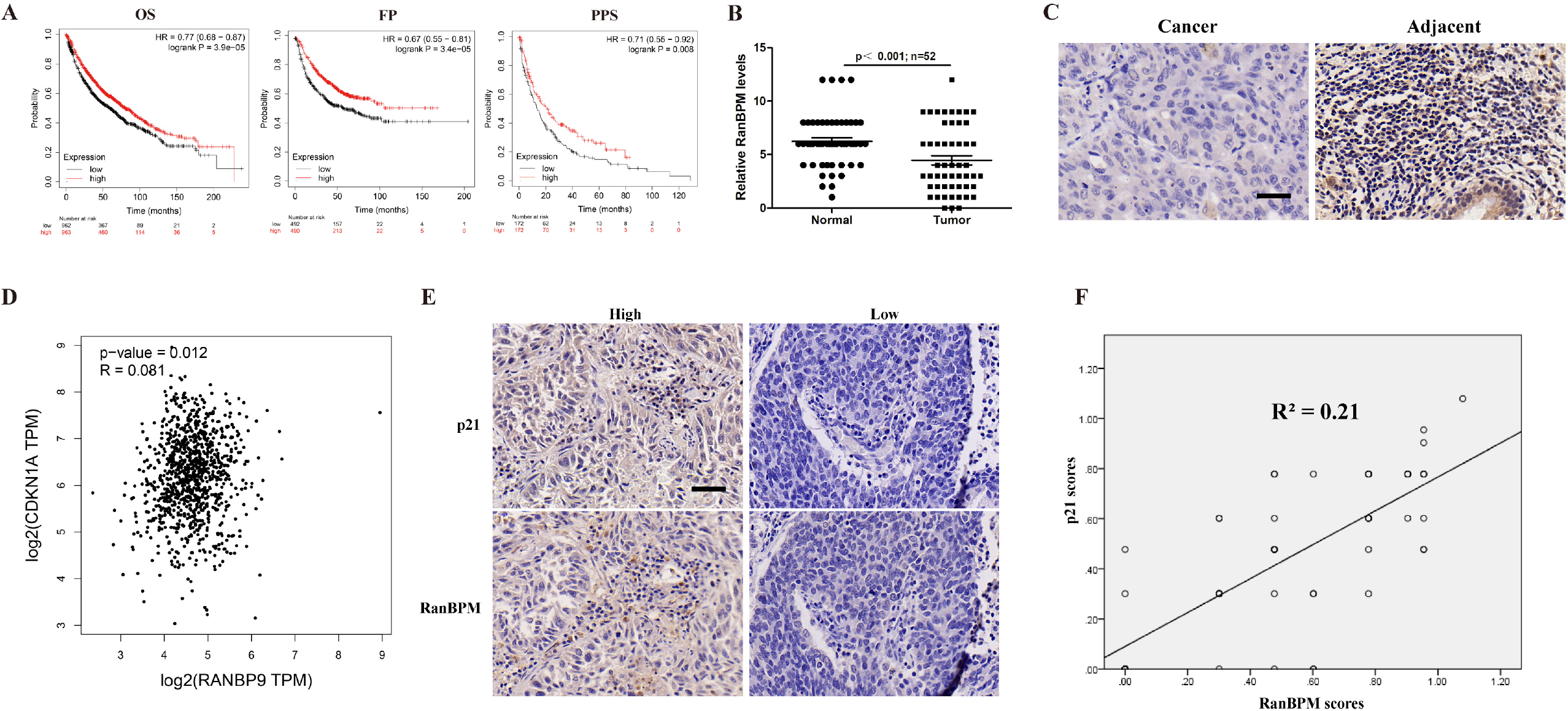
RanBPM is positively correlated with p21, and negatively correlated with the survival of NSCLC patients. (A) Curves of overall survival (OS), progression-free survival (FP) and post-progression survival (PPS) of NSCLC patients were plotted. (B) Quantitation of RanBPM protein levels from IHC images. (C) Representative immunohistochemical (IHC) images of RanBPM in NSCLC tissues or the matched adjacent tissues. (D) Correlation analysis of the mRNA levels of *RanBPM* and *p21* genes in NSCLC samples from TCGA datasets. (E) Representative IHC images of RanBPM and p21 in NSCLC tissues. (F) Regression analysis comparing RanBPM and p21 expression in NSCLC tissues. n=52.

### RanBPM protein interacts with p21

RanBPM, as a scaffolding protein, is a crucial component of multiple-protein complex that mediate diverse cellular functions by modulating and/or assembling with various kinds of proteins [18]. Given RanBPM is positively correlated with p21, we further tested whether RanBPM protein physically interacts with p21. RanBPM or p21 was separately immunoprecipitated from the lysates of A549 cells, and the protein of p21 or RanBPM was detected by western blotting. As shown in Figure 2A and B, both RanBPM and p21 were detected in their individual immunoprecipitated complexes, but not in the isotype-matched negative control IgG complexes. We also detected the colocalization of RanBPM and p21 proteins in the nucleus (Figure 2C). Furthermore, the plasmids encoding Flag-RanBPM or Myc-p21 were transfected into HEK293T cells, and RanBPM or p21 protein was co-immunoprecipitated (co-IP) with an anti-Flag or anti-Myc antibody. The exogenously expressed RanBPM was pulled down by the ectopically-overexpressed p21, while the exogenously expressed p21 was pulled down by the ectopically-overexpressed RanBPM as well (Figure 2D–E). To further validate whether RanBPM physically interacts with p21 protein, we performed GST-pull down assay by using the purified recombinant His-RanBPM and GST-p21 proteins. The GST-p21, but not the GST control, was able to pull down the His-RanBPM protein under cell-free conditions (Figure 2F), demonstrating a direct protein interaction between RanBPM and p21. Collectively, these results suggested that RanBPM physically interacts with p21 protein *in vivo* and *in vitro*.

**Figure 2.**
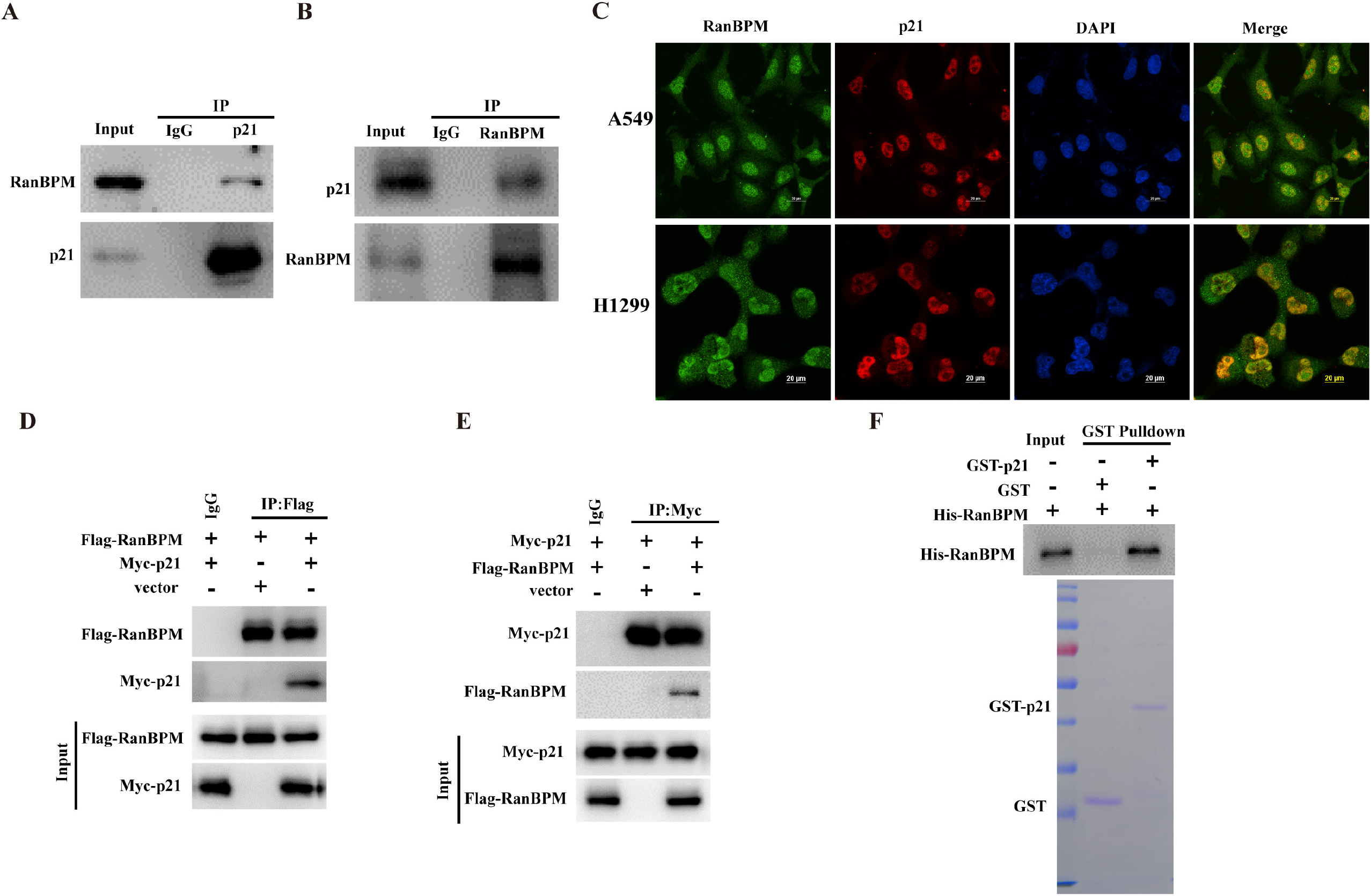
RanBPM protein interacts with p21. (A and B) A549 cell lysates were subjected to immunoprecipitation with isotype control IgG, anti-p21 (A), or anti-RanBPM (B) antibody. The immunoprecipitates were then probed with anti-RanBPM or anti-p21 antibody. (C) The subcellular localization of endogenous RanBPM (green) and p21 (red) in A549 or H1299 cells was visualized using immunofluorescence with anti-RanBPM or anti-p21 antibody. DNA was stained with DAPI, and a merged view of the red and green channels in the same field is shown (merge). (D and E) HEK293T cells were transfected with plasmids encoding Flag-RanBPM and/or Myc-p21. RanBPM or p21 protein was individually immunoprecipitated with anti-Flag or anti-Myc antibody, and RanBPM and p21 was assessed in the immunoprecipitated complex with anti-Flag or anti-Myc antibody. (F) GST, GST-p21, and His-RanBPM produced from bacteria were assessed using western blotting, and the purified RanBPM protein was incubated with GST or GST-p21 coupled to GST-Sepharose. Proteins retained on Sepharose were then blotted with indicated antibodies.

### RanBPM stabilizes and deubiquitinates p21 protein

Since RanBPM protein physically interacts with p21, we next investigated whether RanBPM affects the steady-state levels of p21 protein. As shown in Figure 3A, knockdown of RanBPM with two independent RanBPM specific short interfering RNAs (siRNAs) significantly decreased p21 level in A549 and H1299 cells. Conversely, RanBPM overexpression led to the accumulation of endogenous p21 protein regardless of the p53 status (Figures 3B). To determine whether RanBPM regulates p21 at the level of gene transcription, we measured the mRNA levels of *p21* gene by qRT-PCR after RanBPM downregulation or overexpression. RanBPM did not significantly change the levels of *p21* mRNA (Figure 3C–D). The data suggested that RanBPM regulates the post-translational modifications, rather than gene transcription of *p21* gene.

**Figure 3.**
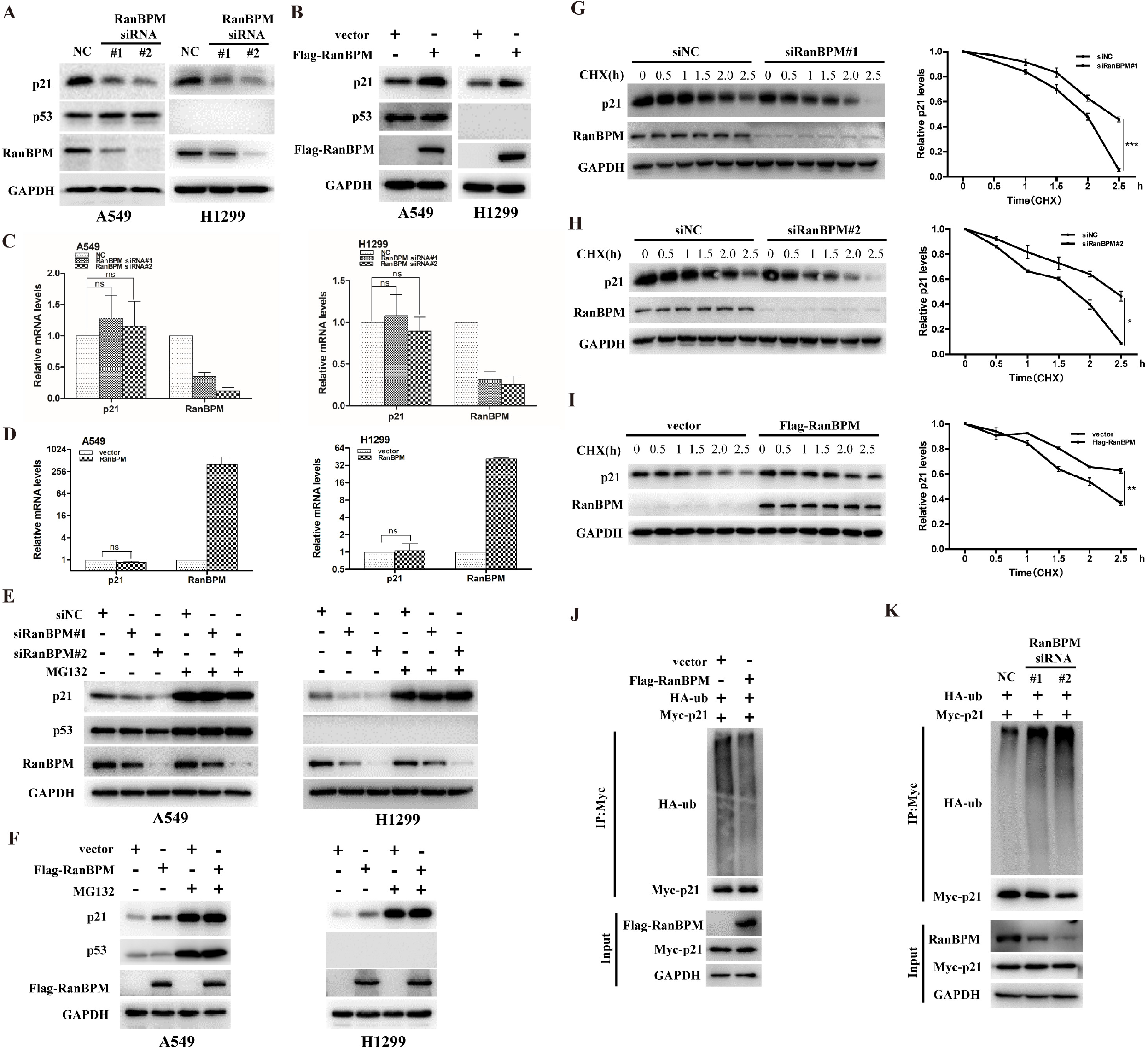
RanBPM stabilizes and deubiquitinates p21 protein. (A) A549 or H1299 cells were transiently transfected with the indicated siRNAs, and the proteins were assessed by western blotting. (B) A549 or H1299 cells were transfected with the indicated plasmids, and the proteins were assessed by western blotting. (C and D) A549 or H1299 cells were infected with the indicated siRNAs (C), or transfected with the indicated plasmids (D), and the mRNA was subjected to qRT-PCR. The error bars represent the SD of triplicate measurements. (E) A549 or H1299 cells were transfected with the indicated siRNAs for 48 h, and then were treated with DMSO or MG132 (20 μM) for additional 6 h. The indicated proteins were analyzed by western blotting. (F) A549 or H1299 cells were transfected with the indicated plasmids for 24 h, and then were treated with DMSO or MG132 (20 μM) for additional 6 h. The indicated proteins were analyzed by western blotting. (G and H) A549 cells were transfected with the siRNAs of scrambled or siRanBPM#1 (G) or siRanBPM#2 (H), and then were treated with 50 μg/mL CHX. The resulting cell extracts were collected at the indicated time points for western blot analysis. The relative values of p21 to GAPDH expression were quantified. (I) A549 cells were transfected with the indicated plasmid constructs, and then were treated with 50 μg/mL CHX. The cells were collected at the indicated times, and proteins were analyzed by western blotting. The relative values of p21 to GAPDH expression were quantified. (J) HEK293T were transfected with plasmids encoding Flag-RanBPM, Myc-p21 and HA-Ubiquitin for 48h, and then treated with MG132 (20 μM) for 6h before harvesting. p21 protein was immunoprecipitated with anti-Myc antibody, and the ubiquitination level of p21 protein was analyzed with anti-HA antibody. (K) HEK293T cells were transfected with the indicated siRNA for 24h, and followed by co-transfection with Myc-p21 and HA-Ubiquitin for another 24h. Cells were treated with MG132 (20 μM) for 6 h before harvesting. p21 protein was immunoprecipitated with anti-Myc antibody, and the p21 protein ubiquitination were tested with anti-HA antibody.

To further elucidate the mechanism by which RanBPM sustains the protein stability of p21, we monitored the protein degradation of p21 when the cells were treated with proteasome inhibitor MG132. In the absence of MG132, RanBPM overexpression or downregulation elevated or declined p21 protein levels in A549 or H1299 cells, while dysregulation of p21 caused by RanBPM could be blocked by the proteasome inhibitor MG132 (Figure 3E–F). The data demonstrated that RanBPM regulates p21 protein by ubiquitin-proteasome pathway. Furthermore, when the protein biosynthesis was inhibited with cycloheximide (CHX), the knockdown of endogenous RanBPM decreased the half-life of the p21 protein (Figure 3G and H), while RanBPM overexpression profoundly extended the half-life of the p21 protein (Figure 3I). To further validate the underlying mechanism by which RanBPM regulates the stability of p21, we measured the levels of polyubiquitination of p21 protein by co-transfecting the plasmids encoding Myc-p21 and HA-Ubiquitin to HEK293T cells. RanBPM overexpression reduced the levels of polyubiquitylated p21 (Figure 3J), whereas RanBPM knockdown significantly increased the levels of p21 polyubiquitylation (Figure 3K).

### RanBPM facilitates p21 deubiquitination in a USP11-dependent manner

It has well known that USP11 plays a key role in the maintenance of p21 protein stability[17]. Interestingly, USP11 recently was identified as a potential binding partner for RanBPM protein [19]. We validated the protein interaction with sequential immunoprecipitation assays. The plasmids encoding Flag-RanBPM, HA-USP11 and Myc-p21 were co-transfected to HEK293T cells, and RanBPM protein was first immunoprecipitated with anti-Flag M2 agarose beads and eluted with Flag peptides. p21 proteins were secondly immunoprecipitated with anti-Myc antibody. The data showed that the USP11 was detected in both immunoprecipitates obtained with anti-Flag or anti-Myc antibodies (Figure 4A). These data indicated that RanBPM, p21 and USP11 were present in the same protein complex.

**Figure 4.**
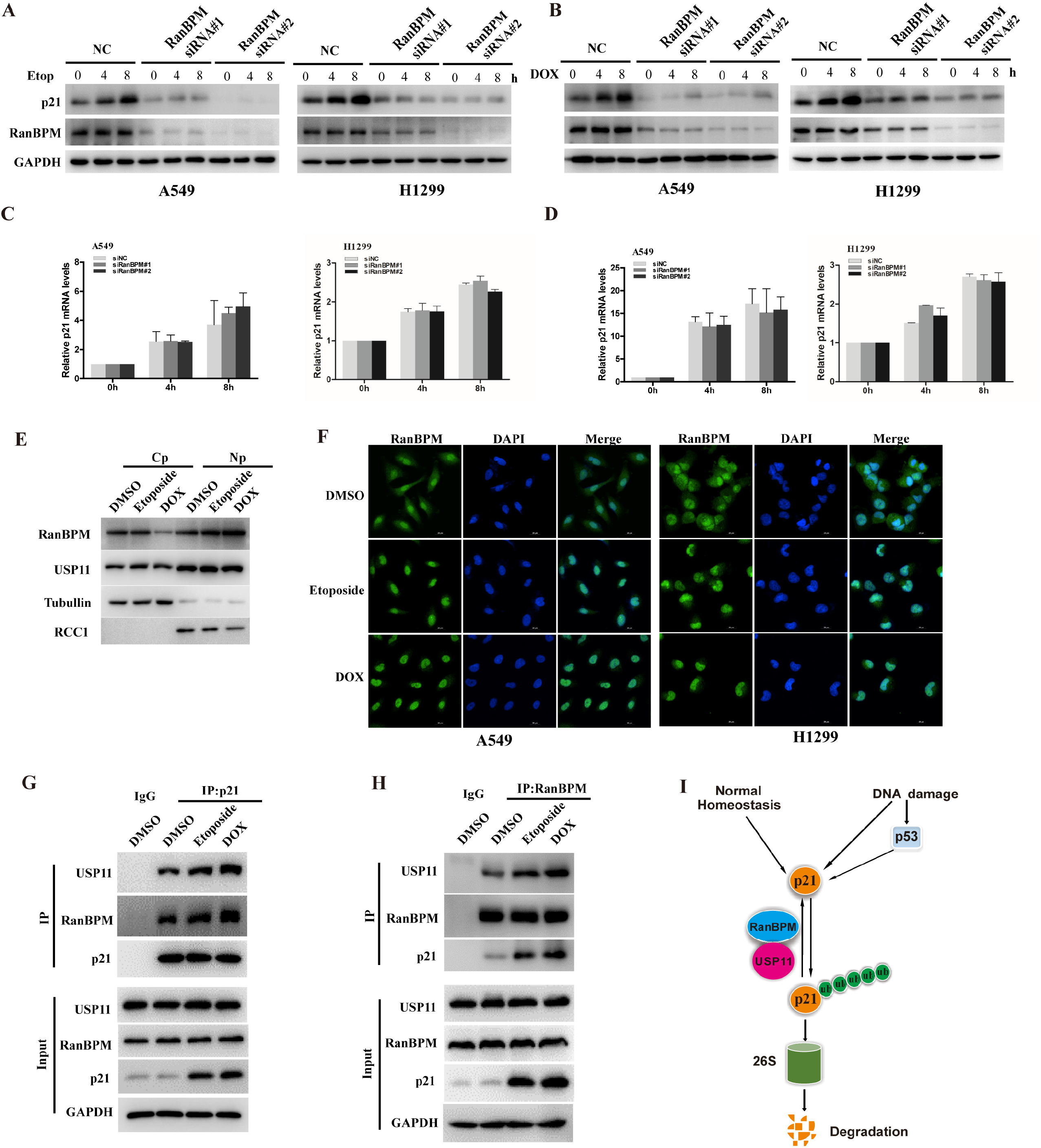
RanBPM facilitates p21 deubiquitination in a USP11-dependent manner. (A) HEK293T cells were co-transfected with plasmids encoding Flag-RanBPM, HA-USP11 and Myc-p21, followed by MG132 treatment. Cell lysates were immunoprecipitated with anti-Flag M2 agarose beads, and eluted with Flag peptides. Eluates were further immunoprecipitated with anti-Myc or control IgG antibodies. Protein samples obtained from each step were analyzed with the indicated antibodies by western blotting. (B and C) A549 cells transfected with the indicated plasmids (B) or siRNAs (C) were treated with MG132 for 6h. Immunoprecipitation were performed with anti-p21 antibody. The immunoprecipitates were then blotted with the indicated antibodies. (D) A549 or H1299 cells were transfected with the indicated siRNAs, and then were transfected with the indicated plasmid constructs. Lysates were blotted with the indicated antibodies. (E) HEK293T cells were co-transfected with the indicated plasmid constructs, and followed by the treatment with MG132 for 6h before harvesting. Cell lysates were immunoprecipitated with anti-Myc antibody, and then were analyzed by immunoblotting with anti-HA antibody. (F) HEK293T cells were transfected with the indicated siRNAs for 24h, and then were transfected with the indicated plasmid constructs. Cell lysates were immunoprecipitated with anti-Myc antibody and analyzed by immunoblotting with anti-HA antibody. (G) A549 and H1299 cells were transfected with the indicated siRNAs, and then were transfected with the indicated plasmid constructs. Lysates were blotted with the indicated antibodies. (H) HEK293T cells were transfected with the indicated siRNAs for 24h, and then were transfected with the indicated plasmid constructs. Cell lysates were immunoprecipitated with anti-Myc antibody and analyzed by immunoblotting with anti-HA antibody.

We further presumed that RanBPM might promote interaction between USP11 and p21, and regulate USP11-dependent deubiquitination of p21. We tested the impact of RanBPM on the protein interaction of USP11 with p21. The ectopic overexpression of RanBPM promoted the protein interaction between USP11 and p21, whereas RanBPM knockdown with siRNAs significantly reduced the USP11-p21 interactions (Figure 4B–C). Furthermore, sub-cellular localization of p21 and USP11 were further detected by immunofluorescent staining after RanBPM was knocked down in A549 cells. The knockdown of RanBPM did not seem to change the sub-cellular localization of p21 and USP11 (Figures S2).

Because RanBPM affected the interaction of USP11 with p21, we hypothesized that RanBPM is involved in USP11-mediated regulation of p21. To address this, Flag-USP11 was transfected into RanBPM-knockdown cells. As expected, the effect of USP11 on p21 markedly decreased after RanBPM knockdown (Figure 4D). Ectopical overexpression of RanBPM dramatically promoted, whereas RanBPM knockdown decreased USP11-mediated p21 deubiquitination (Figure 4E–F). These results suggest that RanBPM plays a vital role in USP11-mediated regulation of p21.

As well known, RanBPM is a scaffold protein without any enzymatic activity[18], we deduced that RanBPM sustains p21 protein stability depending on the deubiquitinase activity of USP11. USP11 knockdown significantly reduced the protein levels of p21, which has been reported in a previous study[17]. We further clarified whether RanBPM facilitates p21 deubiquitination in a USP11-dependent manner. As shown in Figure 4G and 4H, USP11 knockdown impaired RanBPM-sustained p21 protein stability and the deubiquitination level. These results indicated that RanBPM stabilizes and deubiquitinates p21 protein by promoting the binding of USP11 to p21.

### RanBPM regulates DNA damage response in a p21-dependent manner

A549 and H1299 cells with RanBPM knockdown showed a lower ATM activation and defective homology-directed repair (HDR), and DNA damage induced more cell apoptosis [9, 20]. To demonstrate the crucial roles of RanBPM in DDR, we measured the cell proliferation and clonogenic formation in response to genotoxic stress using CCK-8 assays. The data indicated that RanBPM knockdown with siRNAs increased the sensitivity of A549 to genotoxic stress (Figure 5A–C). The rescue of exogenous p21 in the RanBPM-depleted cells fully reversed the effect of RanBPM ablation (Figure 5D–G). These data suggested that the RanBPM-mediated DDR is dependent on p21.

**Figure 5.**
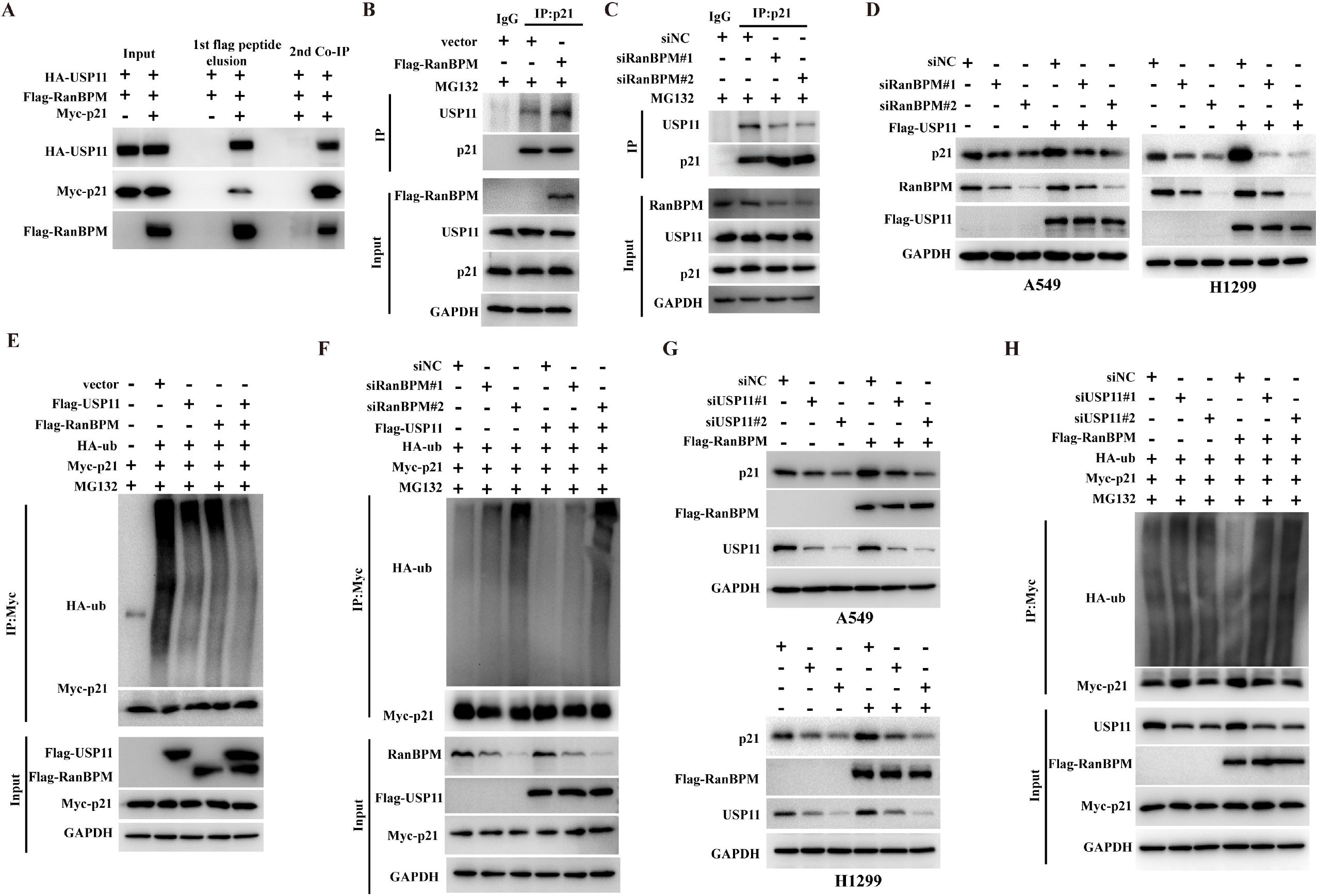
RanBPM regulates DNA damage response in a p21-dependent manner. (A-C) A549 cells were transfected with the indicated siRNAs, and then the cells were treated with 5 μM Etop (B) or 0.5μM Dox (C). (D-G) A549 or H1299 cells were co-transfected with the indicated siRNAs and plasmid constructs, and the cells were treated with 5 μM Etop or 0.5μM Dox. The same number of cells were seeded for colony formation. Data are representative of three independent experiments and values are expressed in mean±SEM (***p<0.001).

### DNA damage promotes the translocation of RanBPM into the nucleus and regulates p21 protein stability

It was reported that p21 regulates DNA damage response (DDR) by p53-dependent or independent pathways[21, 22], Since RanBPM stabilizes and deubiquitinates p21 protein, we next investigated whether DNA damage elevates p21 protein levels through RanBPM-mediated pathways. In agreement with previous reports[23], etoposide elevated p21 protein levels in A549 and H1299 cells (Figure 6A and B), while etoposide-elevated p21 protein levels significantly decreased in the RanBPM-depleted cells (Figure 6A). Furthermore, RanBPM knockdown significantly decreased the doxorubicin-triggered p21 elevation (Figure 6B), but had no impact on the mRNA levels of *p21* gene (Figure 6C and D). These results suggested that RanBPM regulates DNA damage-elevated p21 protein.

**Figure 6.**
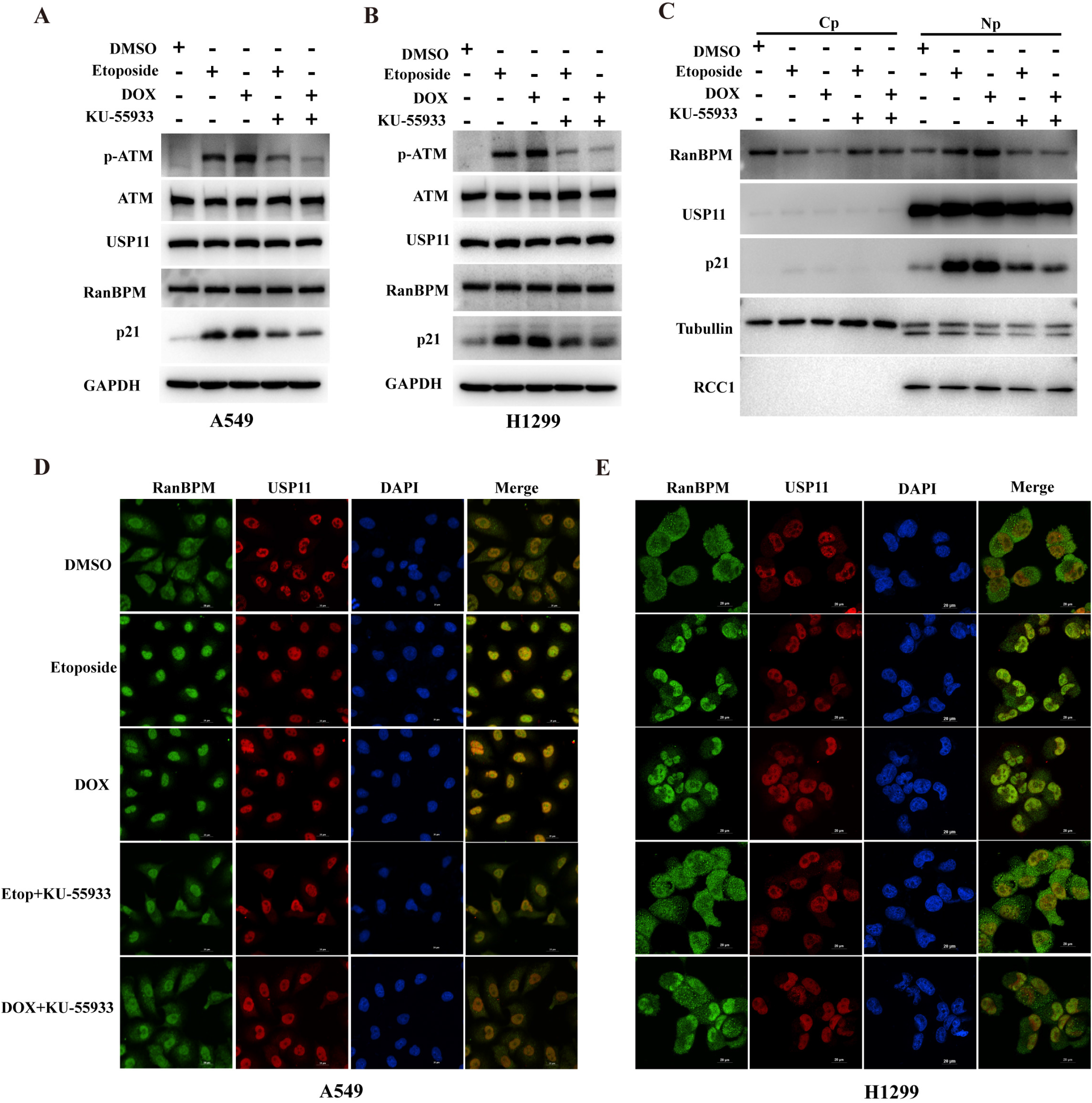
DNA damage promotes the translocation of RanBPM into the nucleus and regulates p21 protein stability. (A-D) A549 or H1299 cells were transfected with the indicated siRNAs, and then were treated with DMSO, 5μM Etoposide (Etop) or 0.5μM doxorubicin (Dox). The cells were collected at the indicated time points, and cell lysates were subjected to western blotting (A and B). The mRNA was subjected to qRT-PCR (C and D). The error bars represent the SD of triplicate measurements. (E) A549 cells were treated with DMSO, 5μM Etop or 0.5μM Dox for 8h. After cell fractionation, the subcellular fractions were blotted with the indicated antibodies. (Cp, cytoplasmic; Np, nuclear). (F) A549 cells and H1299 cells treated DMSO, 5μM Etop or 0.5μM Dox for 8h, the cells were fixed and stained with the fluorescent anti-RanBPM antibodies. DAPI was used for nuclei staining. Scale bars, 20 μm. (G and H) A549 cells were treated with or without 5 μM Etop or 0.5 μM Dox for 8h, and cell lysates were subjected to immunoprecipitation with control IgG, anti-RanBPM or anti-p21 antibody. The immunoprecipitates were then probed with anti-USP11, anti-p21 or anti-RanBPM antibody. (I) Schematic representation of p21 regulated by RanBPM-USP11 axis.

Since p21 acts as a tumor suppressor in the nucleus, we hypothesized RanBPM might translocate into the nucleus to participate in DDR. We performed cell fractionation assays or immunofluorescence to confirm the hypothesis. As shown in Figure 6E, DNA damage significantly elevated the amounts of RanBPM in the nucleus. In addition, DNA damage promoted the translocalization of RanBPM proteins to the nucleus (Figure 6F).

Furthermore, we analyzed the physical protein interaction of RanBPM with p21. A549 cells were treated with etoposide or doxorubicin, and cell lysates were subjected to co-immunoprecipitation with anti-RanBPM or anti-p21 antibody. DNA damage significantly promoted the protein interaction between RanBPM and p21 (Figure 6G and H). These data suggested that RanBPM is indispensable for the p21 protein stability in physiological conditions, or elevated protein levels in response to DNA damage (Figure 6I).

### DNA damage promotes the translocation of RanBPM and regulates p21 protein stability through ATM-mediated pathways

DDR induced posttranslational modifications of proteins such as phosphorylation, which are crucial for controlling protein stability, localization and activity[24]. The DDR signaling pathway orchestrated by the ATM and ATR kinases is the central regulator of this network in response to DNA damage[25]. ATM has been reported as a binding partner of RanBPM[20]. We hypothesized whether DNA damage induced p21 protein accumulation through ATM-mediated translocation of RanBPM. Firstly, etoposide or doxorubicin up-regulated p21 protein levels in A549 and H1299 cells. However, the elevated p21 protein was remarkably decreased by ATM inhibitor Ku55933 (Figure 7A–B). We performed cell fractionation assays to monitor the translocation of RanBPM protein. As shown in Figure 7C, DNA damage significantly increased the amounts of RanBPM proteins in the nucleus. Intriguingly, ATM inhibition by Ku55933 reversed DNA damage-induced nuclear translocation of RanBPM. The data were validated by immunofluorescence (Figure 7D–E). These results suggested that DNA damage significantly promoted the nuclear translocation of RanBPM protein through ATM-dependent pathways.

**Figure 7.**
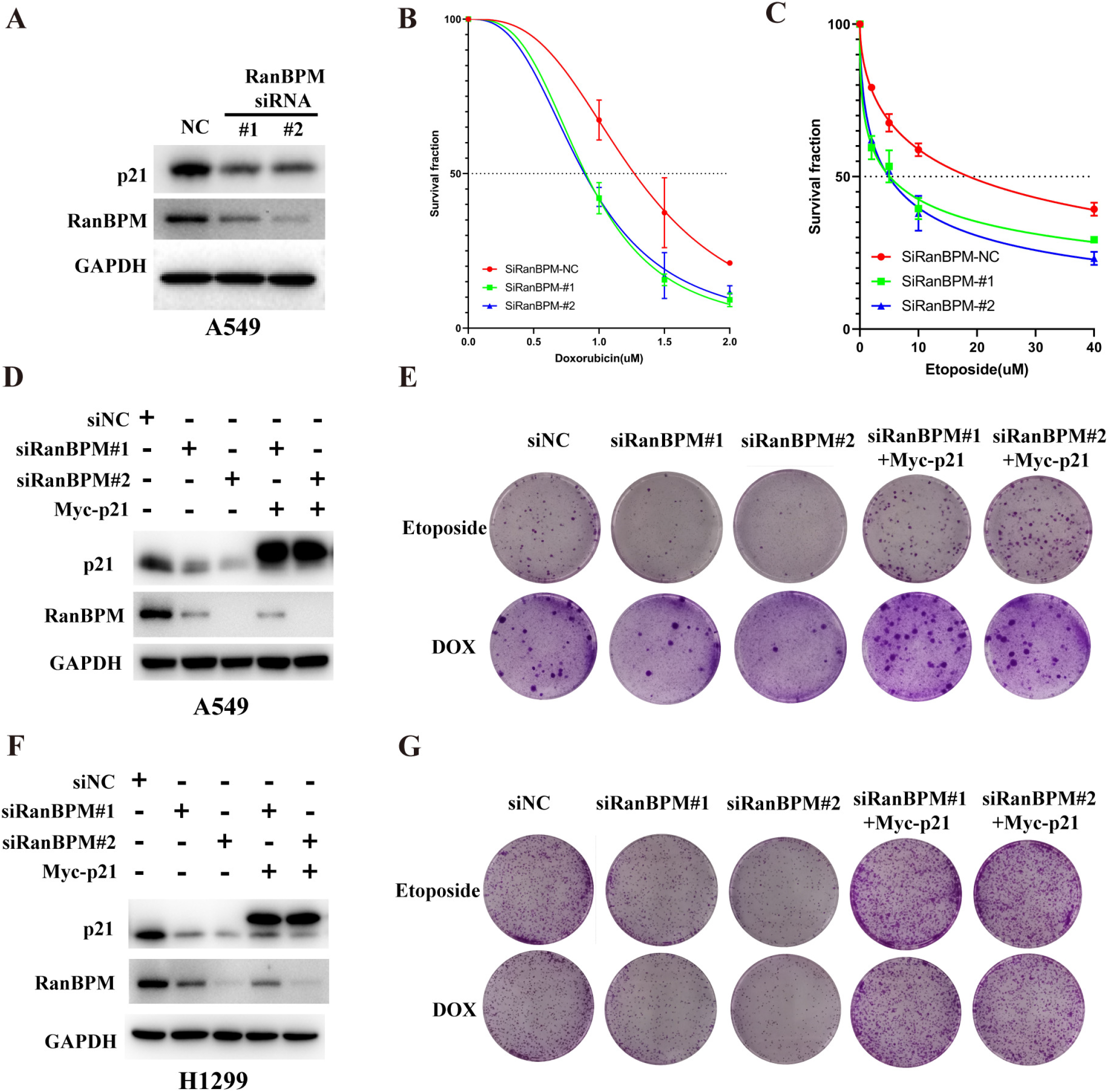
DNA damage promotes the translocation of RanBPM and regulates p21 protein stability through ATM-mediated pathways. (A-C) A549 or H1299 cells were treated with 5μM Etop or 0.5μM Dox in the presence or absence of ATM inhibitor KU-55933 (10μM), and the total proteins were assessed by western blotting. After cell fractionation, the subcellular fractions were blotted with the indicated antibodies(C). (D and E) A549 or H1299 cells were treated with 5μM Etop or 0.5μM Dox in the presence or absence of ATM inhibitor KU-55933 (10μM), and the cells were stained with the indicated fluorescent antibodies. DAPI was used for nuclei staining. Scale bars, 20 μm.

## Discussion

RanBPM as a ubiquitous protein localizes in both the nucleus and cytoplasm, and has emerged as a scaffolding protein to regulate the functions of multiple binding partners by protein-protein interaction. Several studies have reported the vital roles of RanBPM in the regulation of DDR. In the present study, we reported that RanBPM sustains p21 protein stability by tethering a deubiquitinase USP11 to p21, and inhibiting p21 ubiquitination and degradation. Additionally, we also reported that RanBPM is required for DNA damage-induced p21 protein elevation, while the knockdown of RanBPM increases the sensitization of NSCLC cells to DNA damaging agents in a p21-dependent manner (Figure 5E and 5G). Under normal physiological conditions, RanBPM localizes in both the cell nucleus and cytoplasm, while DNA damage promotes the nuclear translocation of RanBPM proteins in a ATM-dependent manner, thereby promoting the interaction of USP11 with p21, and stabilizing p21 protein. These results increased our understanding on the novel roles of RanBPM in the DNA damage elevates p21 protein stability, which is independent of p53 pathway.

p21 is an unstable protein with a relatively short half-life, while the intrinsic and extrinsic stresses such as DNA damage or DNA replication stress rapidly elevated the protein level. The elevation of p21 protein mainly is regulated by post-translational modifications such as phosphorylation or ubiquitylation. Three E3 ubiquitin ligase complexes, SCF^Skp2^, CRL4^Cdt2^, and APC/C^Cdc20^, have been reported to trigger p21 ubiquitylation and degradation at specific stages of the cell cycle. Previous study has reported that USP11 levels varies with cell cycles, and inhibits the ubiquitylation and proteasomal degradation of p21 in a cell-cycle–independent manner. Further studies are needed to fully reveal the detailed mechanism by which RanBPM regulates p21 during cell cycle.

Altogether, the present studies demonstrated that RanBPM plays a crucial role in the USP11-p21 regulatory loop when NSCLC cells were treated by DNA damage agents. DNA damage promotes the nuclear translocation of RanBPM protein, and promotes p21 protein stabilization by facilitating the interaction of p21 with a deubiquitinase USP11 (Figure 8). The project revealed a novel mechanism by which RanBPM regulates p21 protein stability, and plays a critical role in the regulation of DDR, thereby suggesting that it might be a promising therapeutic strategy to develop small compounds destroying the protein interaction between RanBPM and p21.

**Figure 8.**
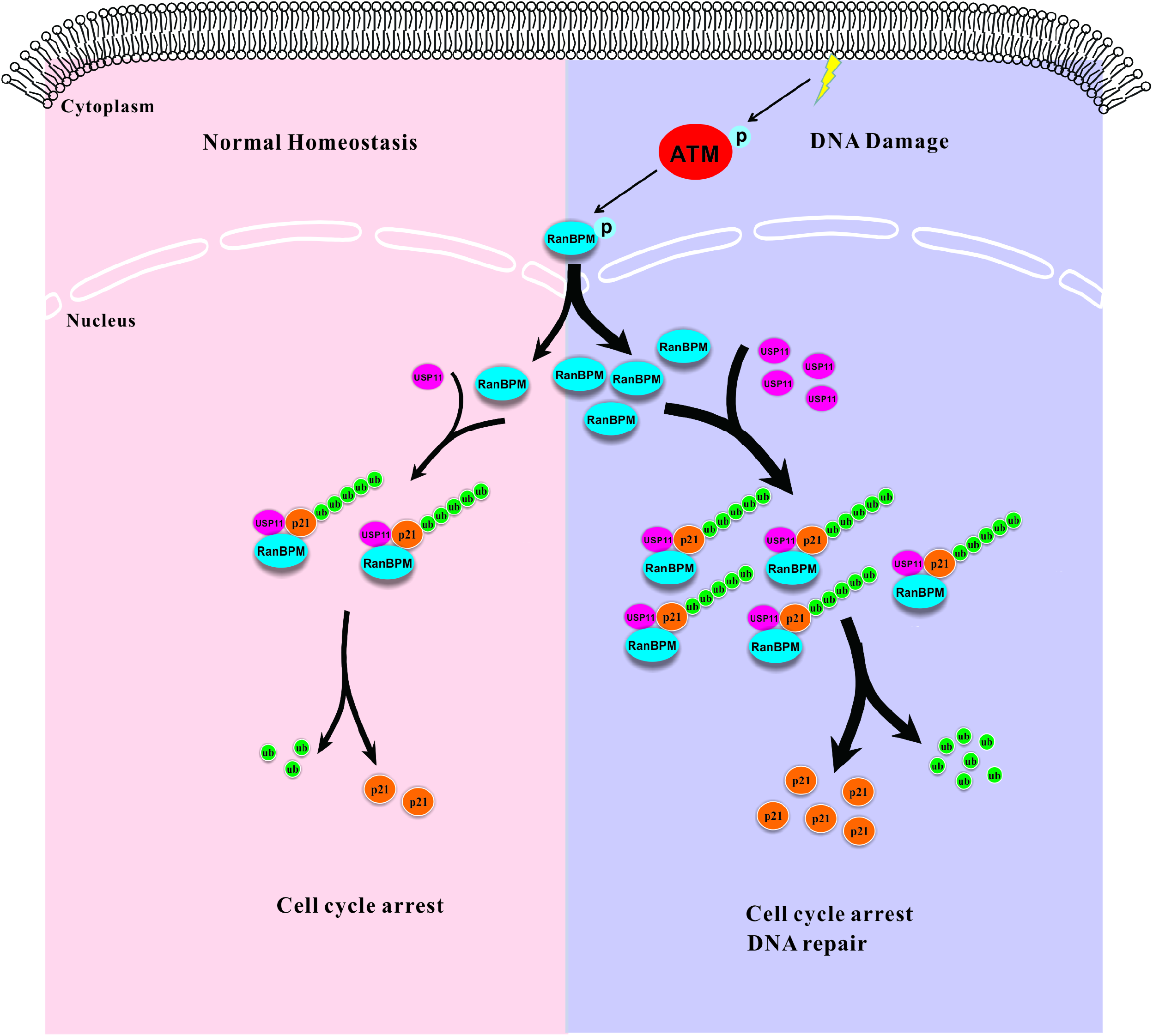
Schematic representation of p21 regulated by RanBPM-USP11 axis.

## Acknowledgements

This study was funded by the National Natural Science Foundation of China (NSFC 82103422, NSFC 81874096), Guangdong Basic and Applied Basic Research Foundation (2021A1515011053), Guangzhou Science and Technology Plan Project (202102020153), National Key Specialty Construction Project of Clinical Pharmacy, High Level Clinical Key Specialty of Clinical Pharmacy in Guangdong Province.

## Author contributions

DTG, XL: data collection and analysis, project development. XYG, ZZB, PYC, YLL, FYM: data collection and analysis. DTG and XL: manuscript writing. LX: Writing-Reviewing and Editing. All authors read and approved the final manuscript.

---

**Figure S1.**
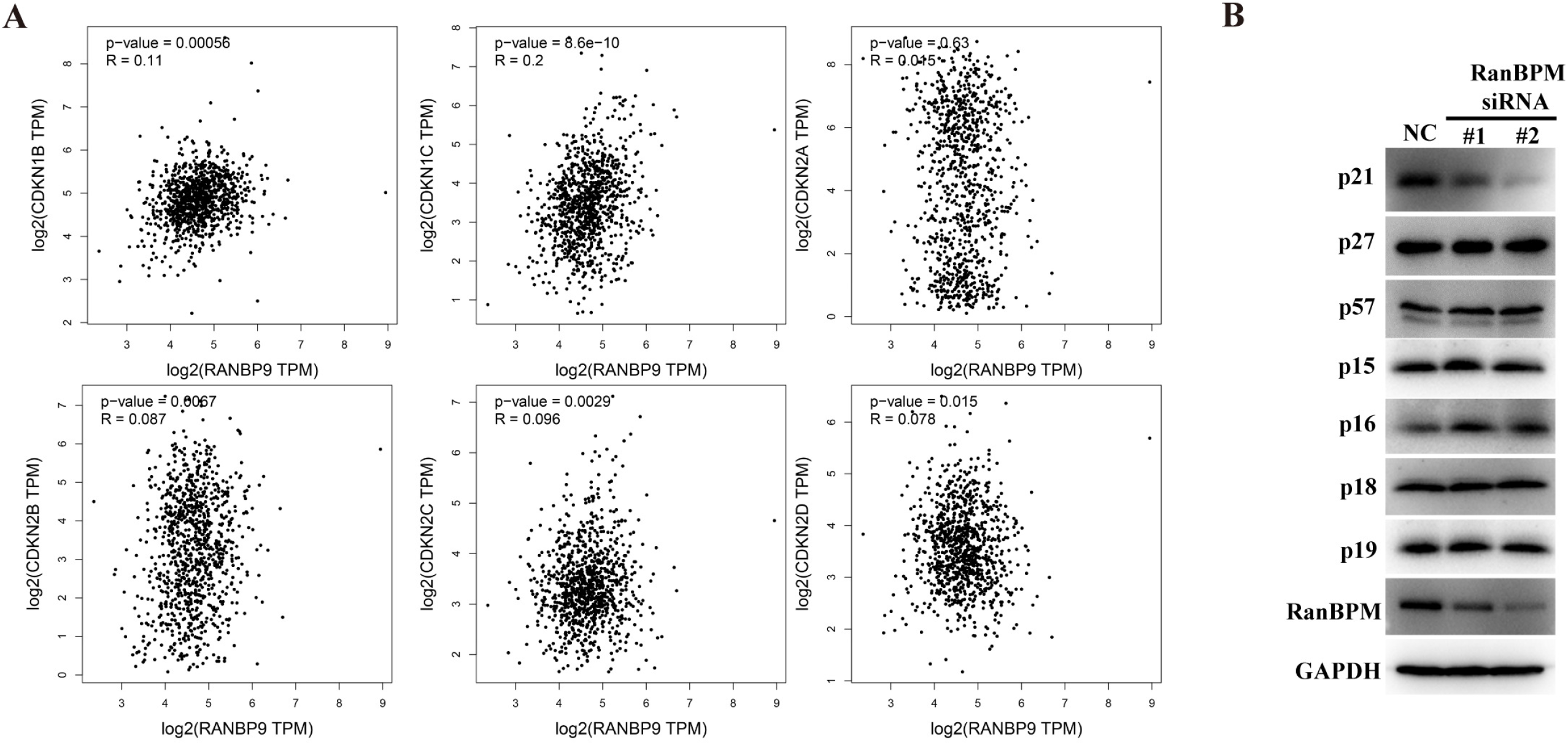

**Figure S2.**
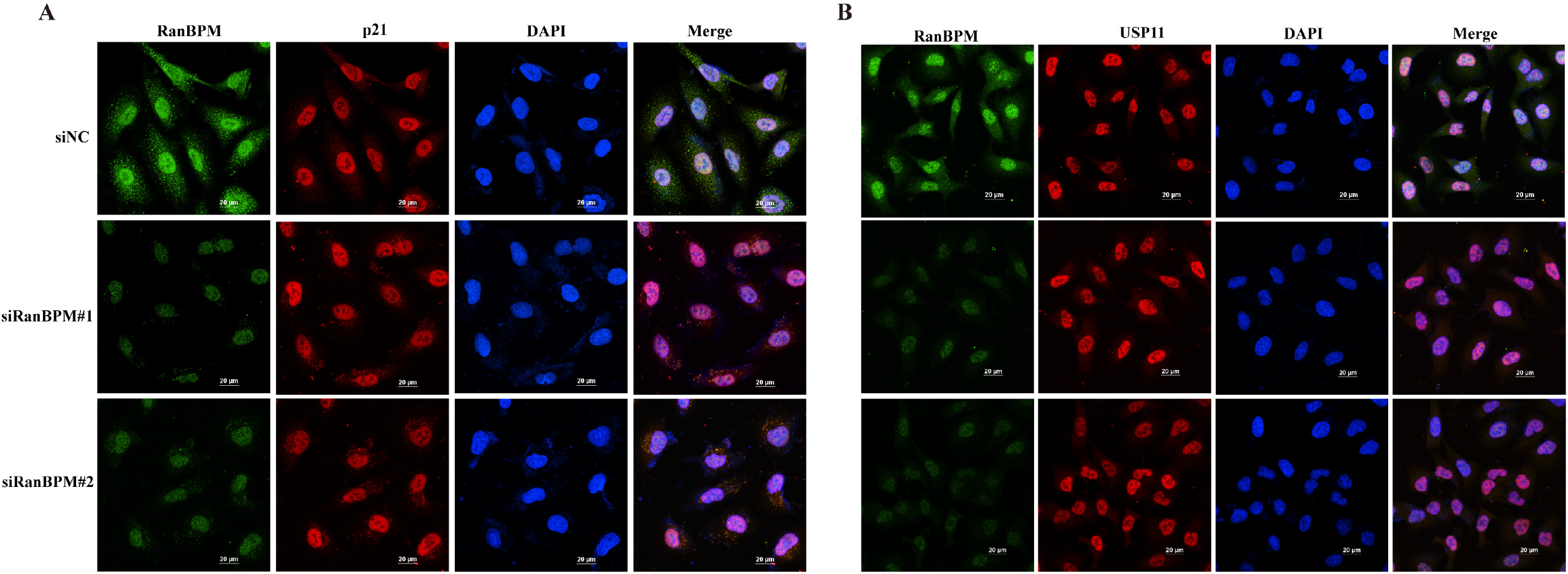

