## supplemental text for "ATM-Mediated Translocation of RanBPM Regulates DNA Damage Response by Stabilizing p21 in Non-Small Cell Lung Cancer Cells"

**Figure S1** (A) Correlation analysis of the mRNA levels of RanBPM and p21 genes in NSCLC samples from TCGA datasets. (B) A549 cells were transiently transfected with the indicated siRNAs, and the proteins were assessed by western blotting.

**Figure S2.** **RanBPM does not regulate the subcellular location of p21 and USP11.** (A and B) A549 cells transfected with the indicated siRNA were treated with MG132 (20 μM) for 6 h, then were fixed and stained. DAPI was used for nuclei staining. Scale bars represent 20 μm.
